# Human stem cell derived neurons and astrocytes to detect novel auto-reactive IgG signature in immune-mediated neurological diseases

**DOI:** 10.1101/2024.02.26.582006

**Authors:** Amandine Mathias, Sylvain Perriot, Samuel Jones, Mathieu Canales, Raphael Bernard-Valnet, Marie Gimenez, Nathan Torcida, Larise Oberholster, Andreas F. Hottinger, Anastasia Zekeridou, Marie Theaudin, Caroline Pot, Renaud Du Pasquier

**Affiliations:** Laboratories of Neuroimmunology, Neuroscience Research Center and Division of Neurology, Department of Clinical Neurosciences, Lausanne University Hospital and Lausanne University, Epalinges, Switzerland; Service of Neurology, Department of Clinical Neurosciences, Lausanne University Hospital and University of Lausanne, Lausanne, Switzerland; Lundin Family Brain Tumor Research Centre, Department of Clinical Neurosciences and Oncology, Lausanne University Hospital and University of Lausanne, Lausanne, Switzerland; Department of Laboratory Medicine and Pathology and Department of Neurology, Center for MS and Autoimmune Neurology, Mayo Clinic, Rochester, MN, United States

**Keywords:** Auto-antibody, human induced pluripotent stem cells, neural cells, NMO seronegative, Auto-immune encephalitis/Paraneoplastic syndrome, Immune-mediated neurological syndromes

## Abstract

**Background and objectives:** Up to 46% of patients with presumed autoimmune limbic encephalitis are seronegative for all currently known CNS antigens. We developed a cell-based assay (CBA) to screen for novel neural antibodies in serum and CSF using neurons and astrocytes derived from human induced pluripotent stem cells (hiPSC).

**Methods:** Human iPSC-derived astrocytes or neurons were incubated with serum/CSF from 99 patients (42 with inflammatory neurological diseases (IND) and 57 with non-IND (NIND)). The IND group included 11 patients with previously established neural antibodies, six with seronegative neuromyelitis optica spectrum disorder (NMOSD), 12 with suspected autoimmune encephalitis/paraneoplastic syndrome (AIE/PNS), and 13 with other IND (OIND). IgG binding to fixed CNS cells was detected using fluorescently-labelled antibodies and analyzed through automated fluorescence measures. IgG neuronal/astrocyte reactivity was further analyzed by flow cytometry. Peripheral blood mononuclear cells (PBMC) were used as CNS-irrelevant control target cells. Reactivity profile was defined as positive using a Robust regression and Outlier removal test with a false discovery rate at 10% following each individual readout.

**Results:** Using our CBA, we detected antibodies recognizing hiPSC-derived neural cells in 19/99 subjects. Antibodies bound specifically to astrocytes in nine cases, to neurons in eight cases and to both cell types in two cases, as confirmed by microscopy single-cell analyses. Highlighting the significance of our novel 96-well CBA assay, neural-specific antibody binding was more frequent in IND (15/42) than in NIND patients (4/57) (Fisher test, *p*=0.0005). Three of three patients with astrocyte-reactive (2 AQP4+ NMO, 1 GFAP astrocytopathy), and 3/4 with intracellular neuron-reactive antibodies (2 Hu+, 1 Ri+ AIE/PNS), as identified in diagnostic laboratories, were also positive with our CBA. Most interestingly, we showed antibody-reactivity in 2/6 seronegative NMOSD, 6/12 probable AIE/PNS, and 1/13 OIND. Flow cytometry using hiPSC-derived CNS cells or PBMC detected antibody binding in 13 versus 0 patients, respectively, establishing the specificity of the detected antibodies for neural tissue.

**Discussion:** Our unique hiPSC-based CBA allows for the screening of novel neuron-/astrocyte-reactive antibodies in patients with suspected immune-mediated neurological syndromes, and negative testing in established routine laboratories, opening new perspectives in establishing early diagnosis of such complex diseases.

## Introduction

The discovery of central nervous system (CNS)-reactive auto-antibodies (Abs) has significantly transformed clinical practice and therapeutic approaches in clinical neurosciences, particularly in the management of neurological disorders such as paraneoplastic syndromes (PNS), autoimmune encephalitis (AIE) or neuromyelitis optica spectrum disorders (NMOSD). Auto-Abs can be identified in diseases with various etiologies including malignancies, auto-immune conditions, and post-infectious syndromes.^1,2^ In the context of suggestive clinical and neuro-radiological features, their precise identification has a profound impact on guiding therapeutic strategies ranging from immunotherapies to cancer therapies.^3,4^ Nevertheless, 7-46 % of the patients with definite or probable autoimmune limbic encephalitis (LE), as defined by 2016 criteria ^5^, still remain seronegative for all currently known neural antigens (Ag) ^5–9^. This suggests that new auto-Abs targets are yet to be discovered.

Similarly, 6-16% of NMOSD patients diagnosed according to the 2015 criteria are seronegative for AQP4 Abs and MOG Abs.^10,11^ In comparison to seropositive cases, in seronegative AIE or NMOSD, the diagnosis is often delayed, hindering appropriate therapeutic measures and impacting outcome. Thus, there is a need for an improved and expedited Ag discovery platform.^8,12–14^

The detection of novel CNS-reactive Abs crucially depends on the antigen source. Preserving the native configuration of the CNS Ag is paramount. Conventional approaches include protein, tissue or cell-based assays. Protein-based assays, such as Western blot, often alter protein configuration, significantly reducing the likelihood of detecting novel antigenic targets for auto-antibodies that depend on conformational structures to bind (e.g. muscle acetylcholine receptor antibodies).^15,16^ Immunofluorescence/immunohistochemistry on animal brain tissues (e.g. rat/ mouse/non-human primate) offers the advantage of preserving some native configuration. However, the use of animal-derived Ags in this assay may prevent the detection of human-specific Abs. Yet human material is scarce (brain biopsies are rarely performed) and available specimens are often in degraded conditions (e.g. autopsy).^17–19^ Last, cell-based assays (CBA) using human CNS primary cells are efficient in binding CNS-specific auto-Abs as the native conformation of human proteins is preserved. Nevertheless, here again, the difficulty to obtain human CNS primary cells represents a real burden in the quest for CNS-specific Abs.^20^

Human iPSC-derived CNS cells emerge as a solution to overcome the limited access to human CNS tissue or CNS primary cells. In this study, we describe a novel CBA using human iPSC-derived neurons and astrocytes as a virtually unlimited source of naturally-expressing neural Ags, to screen for the presence of novel CNS-specific Abs in the serum and cerebrospinal fluid from patients with suspected immune-mediated neurological conditions.

## Materials & Methods

### Study subjects

Patients were enrolled from 2004 to 2022 as part of the ongoing open study COOLIN BRAIN (Cohort Observational Longitudinal Inflammatory Biological Radiological Investigations) aiming at characterizing neuro-immune disorders. Data collection includes clinical, biological, and radiological characteristics. Patients were categorized as follows: 1. AQP4+ and seronegative NMOSD following the 2015 IPND criteria^13^ ; 2. patients diagnosed with definite or suspected AIE/PNS fulfilling diagnostic criteria defined by Graus et al ^5,21^ (autoimmune encephalitis associated or not with a paraneoplastic syndrome; AIE/PNS; 3. other inflammatory neurological disorders (OIND); and 4. non-inflammatory neurological disorders (NIND) as controls (see “results section” for further details. The results of neural/glial-specific auto-Ab testing performed for NMOSD and AIE/PNS patients were collected from reference centers (<u>Labor Krone</u>, Bad Salzuflen, Germany; <u>Clinical Immunology and allergy</u>, Sion, Switzerland; <u>Hospital Clínic of Barcelona</u>, Barcelona, Spain). Patients testing positive for non-CNS-specific auto-Abs in validated diagnostic laboratories at the time of sample collection, such as antinuclear factor (ANA), rheumatoid factor (RF), anti-neutrophilic cytoplasmic Abs (ANCA), anti-double strand DNA Abs (dsDNA) or antiphospholipid Abs were excluded. All samples (serum or CSF) were collected, processed, and stored by the lab through standardized procedures attesting high sample quality (<u>Swiss biobanking platform, Vita label</u>: BBH-NI). This study received approval by our institution’s review board (protocol 2018-01622, COOLIN’BRAIN cohort) and all subjects gave written informed consent before study initiation.

### Human iPSC derived CNS cells: donors, differentiation and culture

Peripheral blood mononuclear cells (PBMC) of two healthy donors were converted into fully characterized hiPSC and neural precursor cells (NPC) as described previously (Age/sex: HD#002: 50/M, cell line ID: LNISi002-B; HD#003: 49/F, cell line ID: LNISi003-A).^22–24^ All donors gave their written informed consent according to regulations established by the responsible ethics committee (CER-VD 2018-01622). Astrocytes were obtained by differentiating NPC in serum free medium following standardized procedure.^22,23^ Media were changed every two to three days and cells were passed when at confluence. Neurons were obtained by lentiviral transduction of lentiviruses encoding for human Neurogenin 2 into hiPSC-derived NPC and induction of expression^25^. Briefly, NPC were plated on poly-L-ornithine/laminin-coated flasks at 50’000 cells/cm^2^ in2µg/ml laminin-supplemented neural expansion medium (DMEM/F-12 + Glutamax, Gibco®; 1x N-2 supplement, Gibco®; 1x B27 supplement without vitamin A, Gibco®; 10 ng/ml FGF-2, PeproTech; 10 ng/ml EGF, Miltenyi). NPC were then transduced with a VSV-G-pseudotyped lentivirus (carrying pCW57.1 plasmid from Addgene # 41393 (gift from David Root) modified to include hNGN2 under the doxycycline promoter). To select for transduced NPC (NGN2-NPC), neural expansion medium was supplemented with 2 ug/mL puromycin after 24 h and changed every 2-3 days. NGN2-NPC were amplified and frozen until use. Neuronal differentiation was induced by culturing NGN2-NPC in neural differentiation medium (DMEM/F-12 + Glutamax, Gibco®; 1x N-2 supplement, Gibco®; 1x B27 supplement without vitamin A, Gibco®) supplemented with doxycycline (2µg/ml). Medium was changed every other day for 6-8 days. All cells were cultivated at 37°C in 5% CO2. Importantly, astrocyte and neuron differentiation profiles were assessed by routine RT-qPCR^22,26^ and immunofluorescence^22,26,27^, confirming the expression of astrocytic or neuronal markers, respectively (See Supplementary Table 1 for primer list, Supplementary Table 2 for the list of reagents used for immunofluorescence stainings, Supplementary Figure 1 for representative results from differentiation profile assessments)

### 96-well hiPSC-derived CNS cell-based assay (96-well hiPSC-derived CNS CBA)

Human iPSC-derived astrocytes and NGN2-NPC were plated on poly-L-ornithine/laminin pre-coated 96-well plates (flat bottom Costar® tissue-treated plates) at 30’000 cells/cm^2^ the day preceding the CBA for astrocytes and at 20’000 cells/well 7-8 days before the CBA for NGN2-NPC to allow in situ neuronal differentiation, respectively.

To detect both extra- and intracellular bound Abs, cells were directly fixed with 4% PFA in PBS for 10 minutes at room temperature (RT) for astrocytes or 100% cold ethanol for neurons for 10 minutes. After washing all wells with PBS, cells were then permeabilized and non-specific sites were blocked using Perm./Block-Buffer (5% NGS + 0.2% saponin in PBS) for 20 minutes at RT. Wells were then washed with PBS and positive controls [anti-CD44 Abs (Miltenyi) for astrocytes; anti-β-tubulin III (Biotechne) Abs for neurons], negative controls (Perm./Block-Buffer only, without serum/CSF nor primary commercial Abs) and patient samples were added into the wells for 35 minutes at RT. Positive controls and serum and CSF samples were diluted in Perm./Block-buffer for extra- and intracellular staining. All serum and CSF samples were run as duplicates and were normalized for their individual IgG concentration (final concentrations tested to minimize unspecific background signal: 15µg/ml for serum and 1.5µg/ml for CSF). Total IgG quantification was determined by the Laboratory of Immunology and Allergy (LIA) of Lausanne, CHUV, Switzerland. Median IgG concentrations were: 10.6 mg/mL in the serum and 32.2 µg/mL in the CSF corresponding to a final median dilution of 1/700 [range: 1/133 – 2540] and 1/22 [range: 1/2; 1/300], for serum and CSF evaluation in our 96-well CBA, respectively.

Biotinylated human anti-IgG secondary Abs also diluted in Perm./Block-buffer were then incubated for 30 minutes at RT in order to detect the presence of bound Abs. After extensive washes in PBS, fluorescent phycoerythrin (PE)-labeled streptavidin was added for another 30 minutes at RT. Negative controls correspond to anti-human IgG detection Abs with PE-labeled streptavidin only. Finally, cells were counterstained with 4’, 6-diamidino-2-phenylindole (DAPI) diluted in PBS for 10 minutes at RT.

Both controls and samples were run in duplicates. Overall IgG binding profile was automatically assessed by measuring PE fluorescence intensity (see section Automated IgG binding profile detection of 96-well hiPSC-derived CNS CBA plates). IgG cellular distribution was further characterized by fluorescence microscopy observation (see section IgG binding profile characterization by fluorescence microscopy observation of 96-well hiPSC-derived CNS CBA plates).

### Automated IgG binding profile detection of 96-well hiPSC-derived CNS CBA plates

Detection of fluorescently labelled cells (fluorescence intensity, FI) was automated using a Synergy® H1 (BioTek®) hybrid multimode plate reader and results were analyzed with Gen5 software (BioTek®, v.3.03.14). To take into account the variable amount of cells at the bottom of each well, raw FI were normalized, using GraphPad Prism® 9 software (Version 9.5.1(733)) in accordance with the following formula: IgG index = [R*aw FI* PE *(well)* − *Mean FI* PE *(negative controls)]/ FI*DAPI *(well).* All wells with restricted number of cells (DAPI signal <2’500) or excessive amount of cells (DAPI signal >20’000) were excluded due to possible bias in IgG index calculation. If the coefficient of variation of the duplicates (CV = standard deviation of duplicate/mean of duplicate*100) was above 50%, the duplicate was considered as discrepant and excluded to ensure robustness of subsequent analyses. A *z-score* of each validated IgG index was then calculated for comparison purposes.

### IgG binding profile characterization by fluorescence microscopy observation of 96-well hiPSC-derived CNS CBA plates

Fluorescence microscopy observations of the 96-well hiPSC-derived CNS CBA was automatized using a high performance EVOS M7000. Four images were acquired per well. Post-image processing was performed using ZEN 2.3 (Zeiss®, v1.3), ImageJ (NIH, v.1.50b) and CellProfiler (version 4.2.1) softwares. Image acquisitions were accomplished using identical parameters for specific Abs, negative controls and patient serum and CSF samples. Images were post-processed identically between all serum samples vs all CSF samples, background levels being specifically adjusted for serum vs CSF. Single cell IgG-associated fluorescence was measured in each individual pictures using CellProfiler (version 4.2.1)^28^ and Cell Profiler analyst (version 3.0.4)^29^ softwares. Mean single cell FI and *z-score* per well were then calculated and strongly correlated with automated plate reader readouts (IgG index [*z-score*]) (Supplementary Figure 2).

### Flow cytometry analyses

Human iPSC-derived astrocytes and neurons were detached and stained as described previously ^22,27,30^. Briefly, detached CNS cells were first stained with Aqua Live/Dead (Life Technologies) for 20 minutes at 4°C. After fixation and permeabilization in Cytofix/Cytoperm buffer (BD Biosciences), cells were washed with PermWash buffer 1x (BD Biosciences) and stained for 35 minutes at 4°C with serum or CSF. All serum and CSF samples were normalized for their IgG concentration (final concentration tested 15µg/ml for serum and 1.5µg/ml for CSF). Cells were then washed and stained with biotinylated human anti-IgG secondary Abs for 30 minutes at 4°C followed by PE-labelled streptavidin (same concentrations as for 96-well hiPSC-derived CNS CBA).

To ensure that IgG reactivity was restricted to CNS cells, CD3+ T lymphocytes were used as an irrelevant source of primary cells. Briefly, a buffy coat of a healthy donor was obtained from the Lausanne interregional transfusion center and total PBMC were isolated as described previously ^31^. PBMCs were then exposed to serum and CSF samples and stained exactly as done for CNS cells except for the addition of an anti-CD3 APC-H7 Ab (BD Biosciences) during the Aqua Live/Dead staining step. Data were acquired on a LSRII flow cytometer (BD Biosciences) and analyzed with FlowJo software (see Appendix I for gating strategies, version 9.1.11, Treestar) (See Supplementary Figure 3 for gating strategies). IgG median fluorescence Intensity (MFI) was used to quantify IgG binding and *z-score* per sample were then calculated for comparison purposes.

### Tissue indirect immunofluorescence (TIF)

All serum and/or CSF samples with IgG staining in neurons or astrocytes using our novel hiPSC-derived CNS cell CBA (Seronegative NMOSD, Suspected AIE/PNS, OIND, NIND) were evaluated by indirect immunofluorescence on a mouse tissue composite (TIF) including brain, kidney and enteric neurons and gut, at the Mayo Clinic Neuroimmunology Laboratory, as previously described.^32^

### Statistical analyses and graphical representations

Both data analysis and creation of figures were performed on R software version 4.3.1 (2023-06-16 ucrt) including specific packages (ggplot2_3.4.3, ggpubr_0.6.0, rstatix_0.7.2, corrplot_0.92). Serum and CSF outliers with a high probability of an association with CNS-reactive Abs were identified using a robust regression and outlier removal test (ROUT test) with a false discovery rate (FDR) at 10 % on calculated IgG index (*z-score*) or IgG MFI (*z-score*). Comparisons of outlier frequencies from one to another group was tested using a two-sided Fisher exact test (p value: *pF* in the text). Correlations between two readouts (96-well plate reader vs microscopy or 96-well plate reader vs flow cytometry) were evaluated using a Spearman’s rank correlation test (p value: *pS* in the text). For all statistical analyses, a p-value inferior to 0.05 was considered significant.

### Data availability

Anonymized data not published within this article will be made available upon reasonable request from a qualified investigator.

## Results

### Study cohort

One hundred and thirty patients were initially enrolled from 2004 to 2022 (Figure 1). Inclusion criteria encompassed patients with an established diagnosis of definite auto-Ab related neurological condition; patients with a high probability of suffering from an Ab-related neurological condition but negative for all CNS auto-Abs tested in validated diagnostic laboratories; OIND patients; and NIND patients. Thirty-one patients were positive to non-CNS-specific auto-Abs and were thus excluded from further analyses. Therefore, 99 patients were finally studied. This cohort included 57 NIND control patients and 42 patients with inflammatory neurological diseases (IND). The latter category included four definite AQP4+ NMO; six seronegative NMOSD; seven definite autoimmune neurological disorders (autoimmune encephalitis associated or not with a paraneoplastic syndrome; AIE/PNS); 12 suspected AIE/PNS; and 13 OIND (Figure 1, Table 1 and Supplementary tables 3-7). To screen for the presence of CNS-specific Abs in selected patients, hiPSC-derived astrocytes (Figure 2) and neurons (Figure 3) were seeded in 96-well plates and exposed to serum and CSF from the 99 study participants.

**Figure 1.**
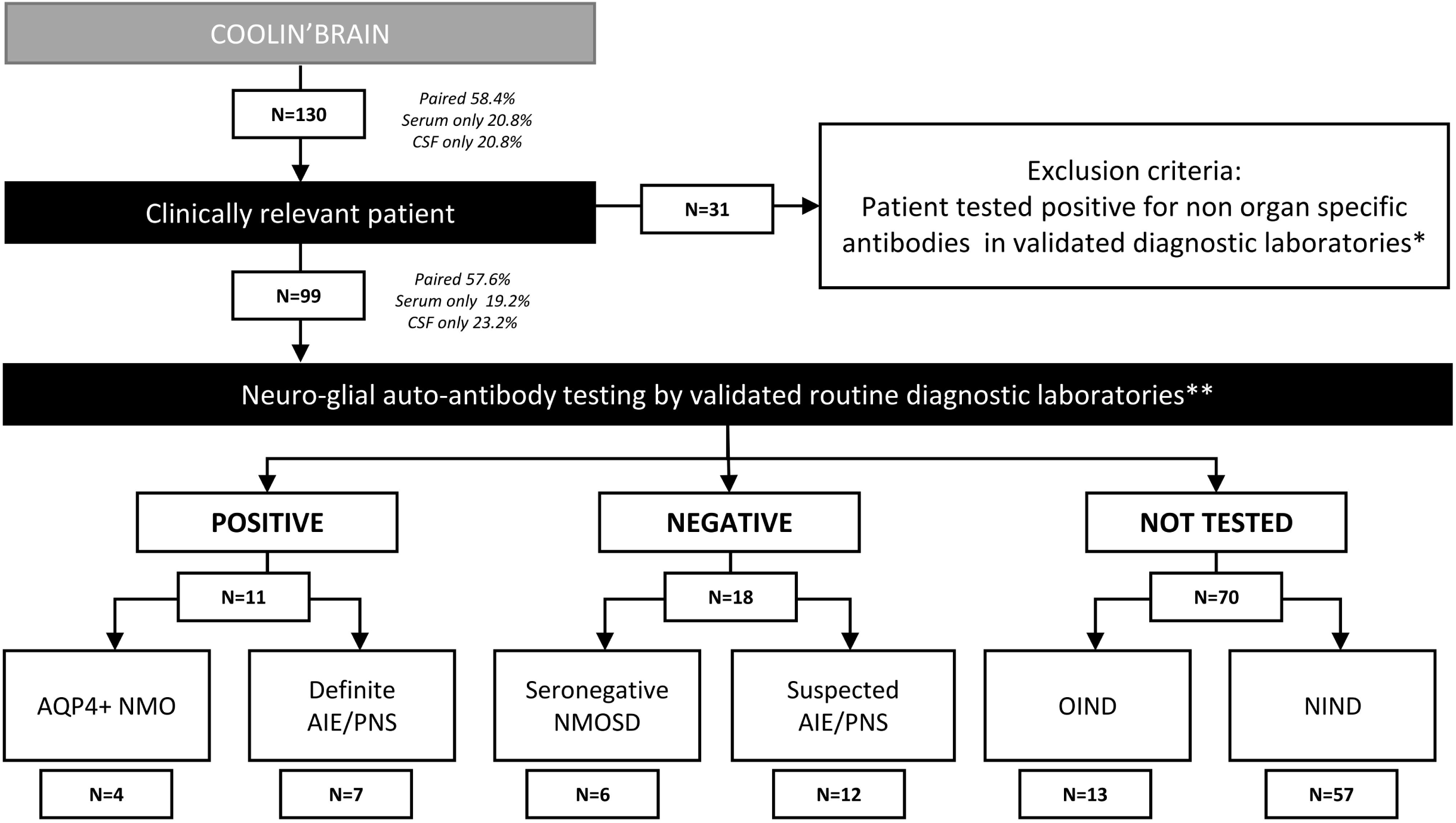
Flow chart of the study design followed for patient recruitment. AIE: autoimmune encephalitis; PNS: paraneoplastic syndrome; NMOSD: neuromyelitis optica spectrum disorder * Non-organ specific antibodies for example ANA, ANCA, dsDNA, Jo, SSA, SSB ** Neural/Glial specific auto-antibody testing routinely included in evaluation panels for suspected autoimmune and paraneoplastic disorders

**Figure 2:**
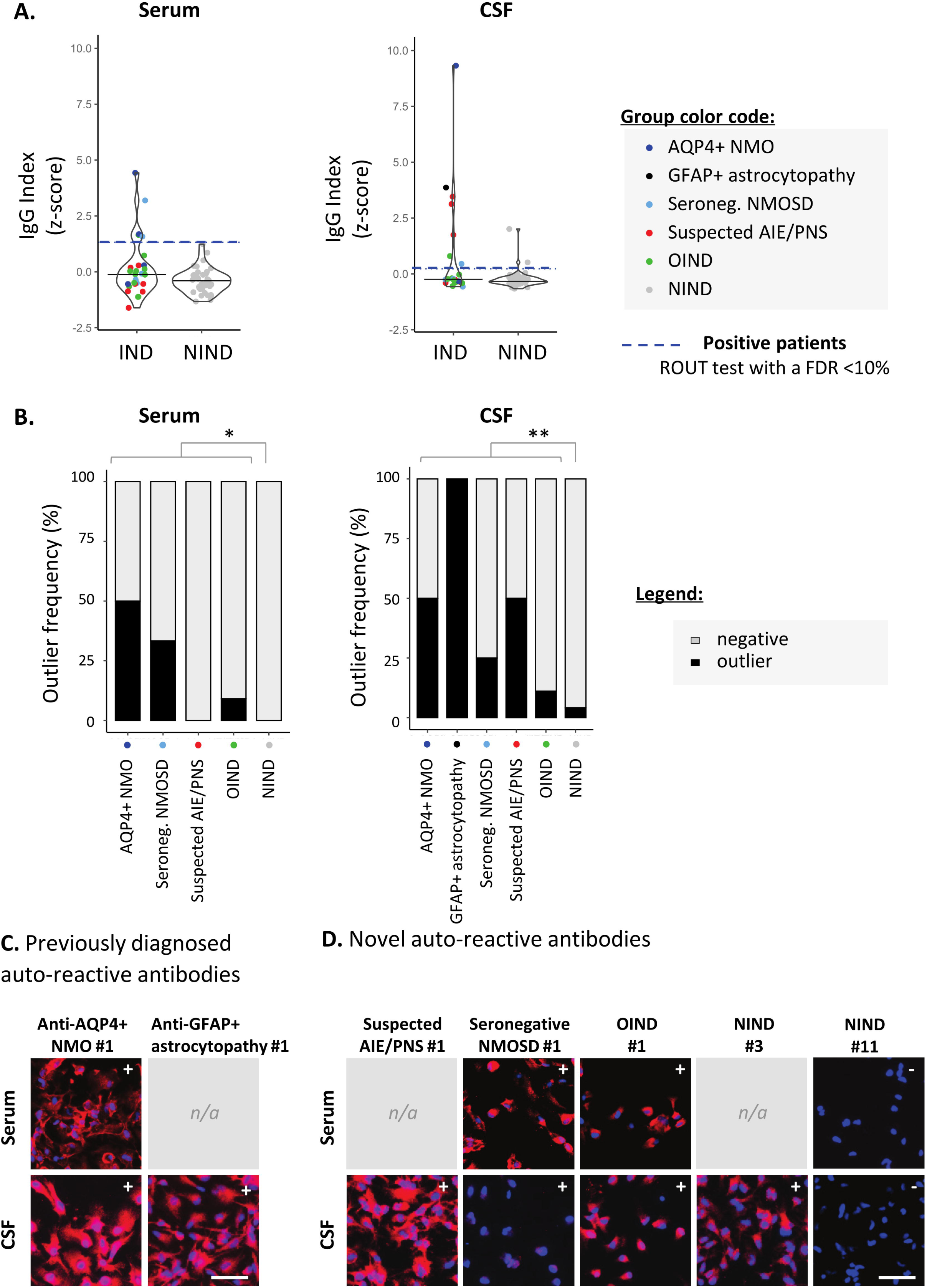
Detection of astrocyte-reactive IgG in sera and CSF of patients with a novel 96-well hiPSC- derived astrocyte cell-based assay (CBA). Human iPSC derived astrocytes were used to screen for CNS-reactive Abs associated with inflammatory and non-inflammatory neurological syndromes (IND, NIND, respectively). Cells were exposed to selected serum and CSF (Figure1). IgG-bound to astrocytes were detected in an intracellular 96-well CBA. **(A)** FI were assessed using a Synergy^®^ microplate reader (left, serum; right, CSF). IgG indexes (*z-score*) are represented (see Methods for details). Each dot represents the mean of duplicates. The black plain lines represent the median of the group. Each clinical subgroups are represented in different colors. The blue dotted line represents the statistically-determined limit above which outliers (defined as positive samples) were identified using a ROUT test with an FDR at <10% (see legend for color/group correspondence). Noteworthy, the patient suffering from GFAP+ astrocytopathy (CSF only) is represented in black as the only definite AIE/PNS patient with reported astrocyte-reactivity. **(B)** The frequency of outliers in each clinical subgroups are represented in black as bar graphs in the serum (left panel) or in the CSF (right panel). Differences in outlier frequencies between IND and NIND groups were tested using Fisher test: *, *p<*0.05; **, *p<*0.01. **(C-D)** Wells of the astrocyte-based CBA were observed by fluorescence microscopy. Representative microscopic observations are depicted for an AQP4+ NMO patient and a patient suffering from GFAP+ astrocytopathy tested positive by validated diagnostic laboratories (C) and for patients tested negative or not tested by validated diagnostic laboratories (D). The presence of IgG bound to astrocytes are detected in red, nuclei staining (DAPI) appears in blue. Symbols (+ or -) depicted in the upper right corner represent the results of CBA either positive outlier (+) or non-outliers (-) sample. White bars represent 50μm. n/a; no paired sample available.

**Figure 3:**
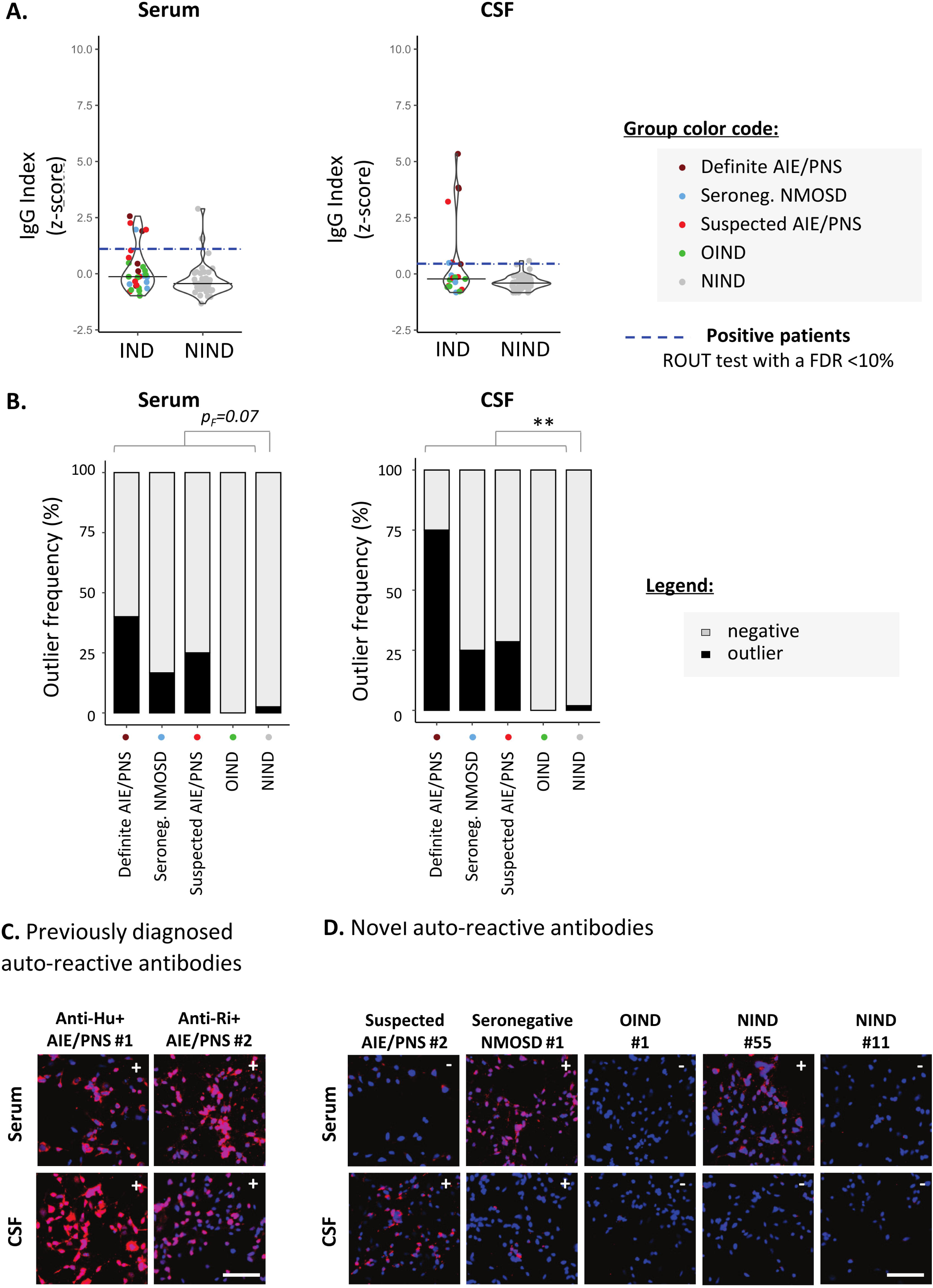
Detection of neuron-reactive IgG in sera and CSF of patients with a novel 96-well hiPSC- derived neuron cell-based assay (CBA). Human iPSC derived neurons were used to screen for CNS-reactive Abs associated with inflammatory and non-inflammatory neurological syndromes (IND, NIND, respectively). Cells were exposed to selected serum and CSF (Figure 1). IgG-bound to neurons were detected in an intracellular 96-well CBA **(A)** FI were assessed using a Synergy microplate reader (left, serum; right, CSF). IgG indexes (*z-score*) are represented (see Methods for details). Each dot represents the mean of duplicates. The black plain lines represent the median of the group. Each clinical subgroups are represented in different colors. The blue dotted line represents the statistically-determined limit above which outliers (defined as positive samples) were identified using a ROUT test with an FDR at <10% (see legend for color/group correspondence). **(B)** The frequency of outliers in each clinical subgroups are represented in black as bar graphs in the serum (left panel) or in the CSF (right panel). Differences in outlier frequencies between IND and NIND groups were tested using Fisher test *pF*: ns, *p>*0.05; **, *p<*0.01. **(C-D)** Wells of the neuron-based CBA were observed by fluorescence microscopy. Representative microscopic observation are depicted for anti-Hu+ or anti- Ri+ PNS patients tested positive by validated diagnostic laboratories (C) and for patients tested negative or not tested by validated diagnostic laboratories (D). The presence of IgG bound to neurons are detected in red, nuclei staining (DAPI) appears in blue. Symbols (+ or -) depicted in the upper right corner represent the results of CBA either positive outlier (+) or non-outliers (-) sample. White bars represent 50μm. n/a; no paired sample available.

**Table 1.** Clinical data of the 99 patients enrolled in this study.

| Group | Diagnostic | Number | M/F ratio | Median age<br>[IQR] (years) | Median Disease duration<br>[IQR] (years) |
| --- | --- | --- | --- | --- | --- |
| **IND** | AQP4+ NMO<sup>a</sup> | 4 | 0/4 | 36.3 [18.3] | 7.04 [18.58] |
|  | Seronegative NMOSD<sup>b</sup> | 6 | 1/5 | 34.9 [9.8] | 1.09 [6.23] |
|  | Definite AIE/PNS<sup>c</sup> | 7 | 2/5 | 59.7 [18.7] | 0.11 [0.39] |
|  | Suspected AIE/PNS<sup>d</sup> | 12 | 7/5 | 66.3 [32.9] | 0.18 [0.267] |
|  | OIND<sup>e</sup> | 13 | 5/8 | 40.4 [24.5] | 0.03 [0.54] |
| **NIND<sup>e</sup>** |  | 57 | 17/40 | 45.3 [24.8] | 0.79 [2.63] |
| **Total** |  | **99** | **32/67** | **44.7 [28.9]** | **0.31 [1.98]** |

Notes:
<sup>a</sup> AQP4+ NMO: see Supplementary Table 3 for subject details.
<sup>b</sup> Seronegative NMOSD: see Supplementary Table 4 for subject details.
<sup>c</sup> Definite PNS/AIE: see Supplementary Table 5 for subject details.
<sup>d</sup> Suspected AIE/PNS: see Supplementary Table 6 for subject details.
<sup>e</sup> OIND and NIND: see Supplementary Table 7 for subject details.
IND : inflammatory neurological diseases; PNS : paraneoplastic syndrome; AIE : autoimmune encephalitis; NMOSD : neuromyelitis optica spectrum disorder; MS : multiple sclerosis; OIND : other inflammatory neurological diseases; NIND: non-inflammatory neurological diseases

### Astrocytes reactive-antibodies

Relying on the rapid automated plate reader IgG measurements, we identified astrocyte-specific IgG in in 9/36 (25%) IND vs 2/57 (3.5%) NIND patients (*pF*=0.003; Figure 2 A). Astrocyte-reactive Abs were more frequently found in the CSF only (5/11; 45.4%) than in the serum only (3/11; 27.3%) or in both serum and CSF together (3/11; 27.3%). Validating the ability of our 96-well hiPSC-derived astrocyte-based CBA to detect astrocyte-specific IgG, we observed a strong astrocyte signal in the CSF of a patient with autoimmune GFAP astrocytopathy and in the serum and CSF of two patients with NMOSD and AQP4-Abs in a routine reference laboratory at the time of sample collection. Our astrocyte-based CBA was negative in two patients previously diagnosed with AQP4+ NMOSD but tested negative (AQP4+ NMO #4) or with a low AQP4+ Ab titer (AQP4+ NMO #3; 1/32), a titer far higher than the one established in our assay (final serum concentration 15 µg/mL corresponding to a 1/860 dilution for this patient). Most interestingly, using this novel 96-well hiPSC-derived astrocyte CBA, we were able to detect astrocyte-specific IgG in: 2/6 seronegative NMOSD (33.3%); 3/12 Suspected AIE/PNS (25%); and 1/13 OIND (7.7 %) (Figure 2B and Supplementary Table 3-7 for clinical description). These results were further illustrated and confirmed by fluorescence microscopy observations (Figure 2 C-D).

### Neuron-reactive-antibodies

Similar experiments were conducted using hiPSC-derived neurons. We identified neuron-specific IgG in 2/57 NIND (3.5%) and in 8/37 (21.6%) IND study patients (*pF*=0.013, Figure 3A). Contrasting with astrocyte-reactive IgG, which were detected mostly in the CSF, neuron-specific IgG were found similarly distributed between the two compartments: CSF only in 3/10 (30%) and serum only in 4/10 (40%). Antibodies were detected in both serum and CSF in 3/10 (30%). Such as was done for astrocytes, we enrolled six patients with definite neuron-specific Ab-associated neurological disorders: anti-AK5+ AIE/PNS (n=1); anti-PCA-Tr+ AIE/PNS (n=1); anti-NMDAR+ AIE/PNS (n=1); anti-Hu+ AIE/PNS (n=2); and anti-Ri+ AIE/PNS (n=1). We were able to identify 3/6 AIE/PNS patients (50%), namely the two Hu+ and the Ri+ AIE/PNS patients.

As part of the exploratory cohort, we selected 12 patients with suspected AIE/PNS who tested negative in validated diagnostic laboratories using all known neural/glial specific auto-Ab panels, all patients fulfilling the AIE/PNS criteria.^5,21^ Of those 12 patients, 9 had an underlying oncological condition (Table 1, supplementary table 6 for clinical details). Using this novel 96-well hiPSC-derived neuron CBA, we were able to detect neuron-specific Abs in 4/12 patients suspected AIE/PNS patients (33.3%). Interestingly, we could also detect neuron-specific Abs in 1/6 seronegative NMOSD (16.7%) (Figure 3B, Supplementary Table 4 for clinical description). These results were further illustrated and confirmed by fluorescence microscopy observations (Figure 3 C-D).

Finally, IgG staining patterns of serum/CSF samples from suspected AIE/PNS, seronegative NMOSD, OIND and NIND patients found to be positive in our novel hiPSC-derived CNS CBA were evaluated by indirect immunofluorescence on a mouse tissue composite (TIF). None of these patients demonstrated any pattern specific of to-date reported CNS-reactive auto-Ab, supporting our human based model as a new tool to detect new human CNS-reactive auto-Abs.

### Comparison of 96-well CBA and flow cytometry measurements

To examine how our 96-well hiPSC-derived CNS CBA assessment (plate reader + fluorescence microscopy) would compare to flow cytometry measurements in the detection of CNS-reactive auto-Abs, we assessed the IgG-binding profile on detached hiPSC-derived astrocytes or neurons (Figure 4) or CD3+ T cells (the latter being used as control cells; Figure 5). All three cell types were exposed to the same selected serum and CSF samples and the IgG binding profile was assessed by quantifying IgG MFI. Serum or CSF were defined as positive using a ROUT test with a FDR set at 10% on corresponding IgG MFI (*z-score*).

**Figure 4:**
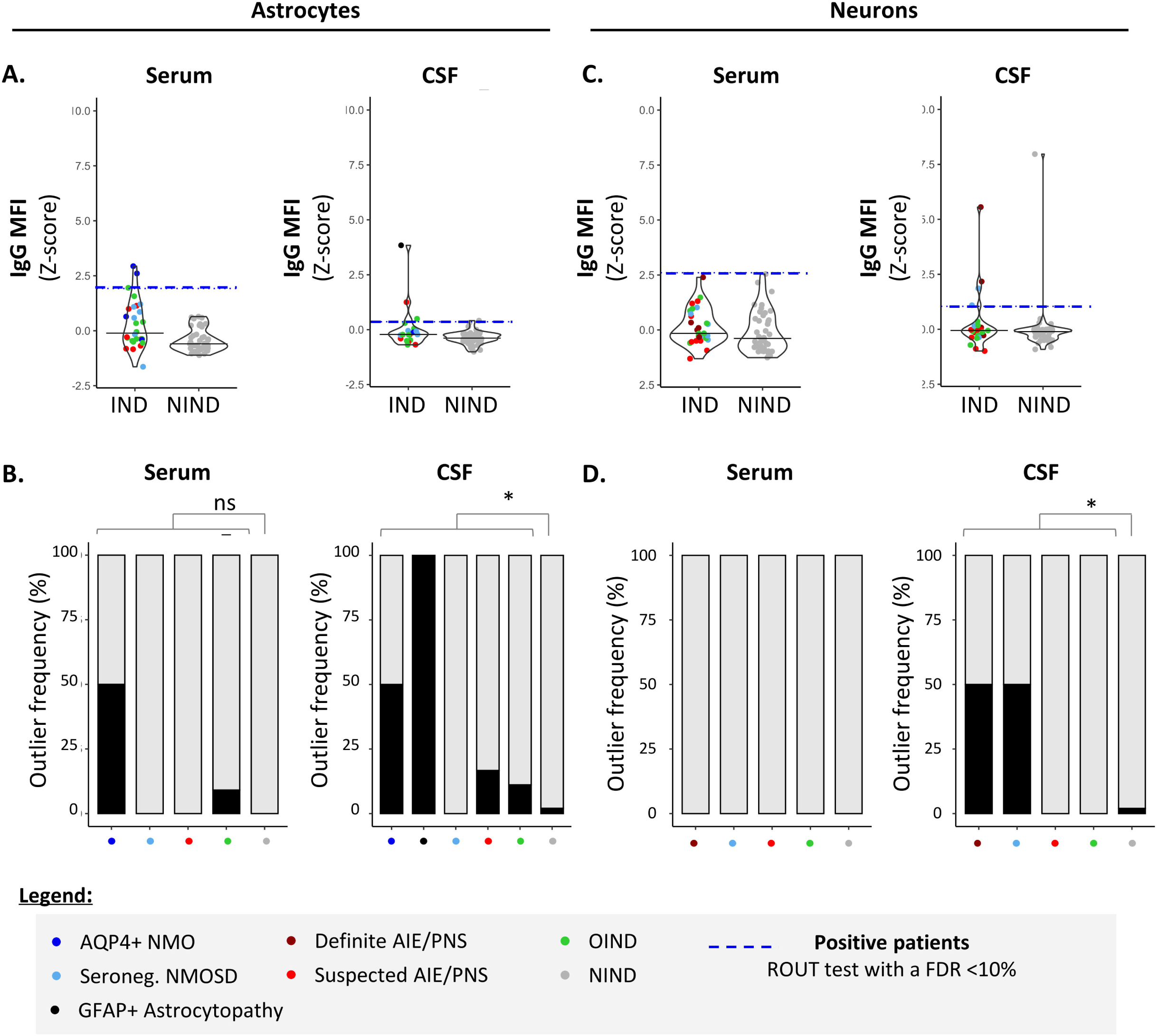
Detection of CNS auto-reactive IgG in sera and CSF of patients by flow cytometry using hiPSC-derived astrocytes and neurons. Human iPSC-derived astrocytes (A-B) or neurons (C-D) were detached and exposed to the exact same serum (left panels) and CSF (right panels) of patients as in Figures 2 and 3. **(A, C)** IgG MFI binding profile (*z-score*) to astrocytes (A) or neurons (C) was quantified by flow cytometry. Each dot corresponds to one sample. The black plain lines represent the median of the group. Each clinical subgroup has a different color. The blue dotted line represents the statistically- determined limit above which outliers (defined as positive samples) were identified using a ROUT test with an FDR at <10% (see legend for color/group correspondence). **(B, D)** The frequency of outliers in each clinical subgroups using either hiPSC-derived astrocytes (B) or neurons (D) are displayed in black as bar graphs in the serum (left panels) or in the CSF (right panels). Differences in outlier frequencies between IND and NIND groups were tested using Fisher test: ns, *p>*0.05; *, *p*<0.05.

**Figure 5.**
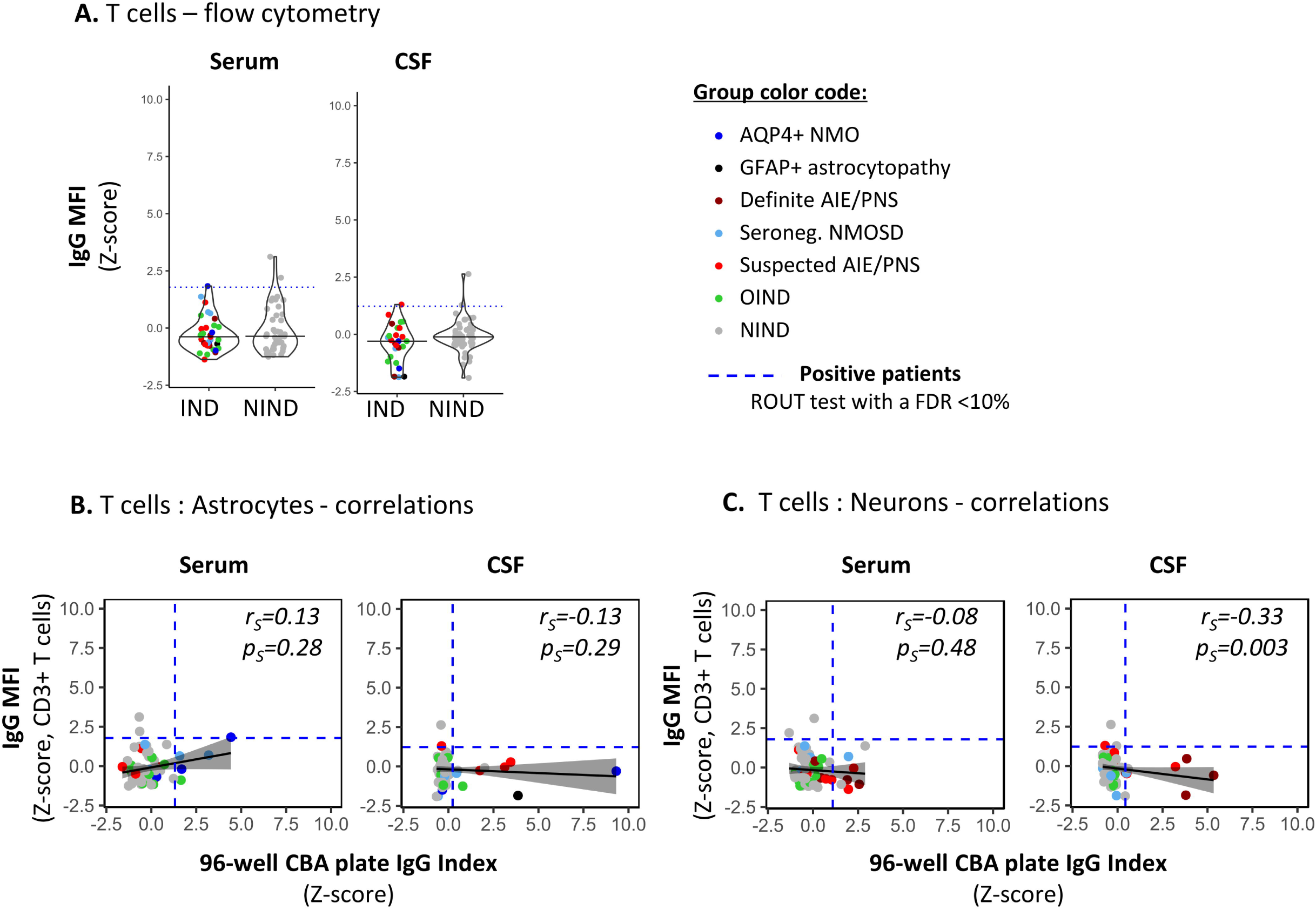
Detection of non-CNS specific auto-reactive IgG in sera and CSF of patients by flow cytometry using irrelevant CD3+ T cells. **(A)** PBMC were exposed to the exact same serum (right panel) and CSF (left panel) as in Figures 2 and 3 for auto-reactive antibody detection (same reagent, same protocol as in Figure 4 for astrocytes and neurons). IgG MFI bound to CD3+ T cells present in PBMC was assessed using flow cytometry. Each dot represents one sample. The black plain lines represent the median of the group. Each clinical subgroups are represented in different colors. The blue dotted line represents the statistically- determined limit above which outliers (defined as positive samples) were identified using a ROUT test with an FDR at <10% (see legend for color/group correspondence). **(B-C)** The absence of positive correlations between IgG index measured by the automated plate reading of 96-well CBA [x-axis] using either hiPSC derived astrocytes (B, Figure2) or neurons (C, Figure3) [x-axis] and CNS-irrelevant CD3+ T cell associated IgG MFI [y-axis] were represented using a generalized linear model (reference line in plain black, confidence interval set at 95% in shadowed area) and tested using a Spearman’s rank correlation test (*rs* and *ps* values on the graphs).

We found that there was a significant correlation between both astrocyte-based readouts i.e. 96-well hiPSC-derived CNS CBA and flow cytometry, but this correlation stood only when astrocytes were exposed to CSF and not to serum (plate reader : IgG index [*z-score*] vs flow cytometry: IgG MFI [*z-score*]) (Spearman correlation statistics: serum, *pS*= 0.15/*rS*=0.18; CSF, *pS*= 0.03/*rS*=0.26). This positive correlation reinforces our hiPSC-derived astrocytes as a reliable source for the detection of astrocyte-specific auto-Abs. However, flow cytometry acquisition tended to be less sensitive than the 96-well hiPSC-derived astrocyte CBA. Comparing flow cytometry versus 96-well hiPSC-derived CNS CBA, we identified 5/34 vs 9/36 IND and 1/57 vs 3/57 NIND, respectively (Figure 4 A-B).

Regarding neurons, we observed no correlation between flow cytometry analyses MFI and 96-well CBA IgG index plate reader (Spearman correlation: serum *pS*=0.37/*rS*=0.11; CSF *pS*=0.08/*rS*=0.20). The less good yield of flow cytometry for this type of CNS cells is likely related to the detachment of hiPSC-derived neurons, a procedure that entails loss of neuronal structure, likely impairing Ag recognition by neuron-reactive auto-Abs. Indeed, comparing neuron-reactive IgG detected by flow cytometry versus 96-well hiPSC-derived CNS CBA, we identified 4/36 vs 8/37 IND and 1/57 vs 3/57 NIND, respectively (Figure 4C-D). These findings align with previous reports indicating that flow cytometry is less sensitive as compared to live or fixed CBA, all of which are techniques commonly used for the diagnosis Ab-related neurological conditions.^33^

Finally, to confirm that serum and/or CSF IgG reactivity as defined by our 96-well hiPSC-derived CNS CBA was specifically directed towards CNS antigens, we assessed the IgG reactivity against primary irrelevant cells (CD3+ T cells) using a flow cytometry assay performed on the same serum/CSF samples (Figure 5). Notably, primary CD3+ T cells are naturally non-adherent cells thus flow cytometry acquisition represents an appropriate method to assess IgG binding profile (Figure 5A). The absence of any correlation between 96-well hiPSC-derived CNS CBA (IgG index (*z-score*)) and IgG reactivity to CD3+ T cells (IgG MFI (*z-score*)) strongly suggests that astrocyte- or neuron-IgG reactivities as identified by the 96-well hiPSC-derived CNS CBA are indeed limited to CNS Ags (Figure 5C-D).

In summary, our novel 96-well hiPSC-derived CNS CBA appears to be a robust tool to detect astrocyte- or neuron-specific Abs in the serum, and especially in the CSF of patients suspected to have neurological syndromes with an autoimmune/paraneoplastic etiology.

## Discussion

Despite indisputable improvements in the diagnosis of immune-mediated neurological disorders facilitated by the detection of CNS-reactive Abs, there is still room for improvement. Indeed, physicians frequently encounter patients suspected of having neurological syndromes with an autoimmune etiology, yet no CNS-specific Abs are identified, even in reference laboratories.^2,3^

In this study, we present a novel platform, namely a 96-well hiPSC-derived astrocyte- and neuron-CBA, enabling the rapid detection of CNS-reactive Abs. This 96-well CBA yielded more robust results as compared to flow cytometry acquisition on detached hiPSC-derived astrocytes or neurons, consistent with other techniques used in validated laboratories.^33^ Using this 96-well CBA to detect CNS-reactive Abs, we identified nine patients with a reactivity profile restricted to astrocytes, eight to neurons, and two to both astrocytes and neurons. However, none exhibited detectable Ab binding to CD3+ T cells (used as non-CNS control cells), suggesting that the observed binding to neurons and/or astrocytes was specific to the neural Ags.

Importantly, we demonstrate that this novel 96-well CBA can replicate findings of established laboratories. More specifically, this assay caught an astrocyte-specific Ab response in 1/1 GFAP astrocytopathy and 2/4 AQP4+ NMOSD (2/2 with positive AQP4-specific humoral immune response demonstrated in established laboratories at the same time as we performed our CBA assay). As per the two AQP4+ patients who tested negative in our 96-well hiPSC derived astrocyte CBA, one was also negative and the other exhibited a low titer (1/32) in a reference lab. Furthermore, both were on immunomodulatory treatment (Mycophenolate Mofetil) for 7.1 years and 13.7 years at sampling date. It is known that AQP4+ positivity can be transitory in NMO, especially upon long-term immunosuppressive treatment. Indeed, over 3.7 years, one third of the treated AQP4+ NMO patients seroreverts to AQP4 negative.^34,35^

The CBA assay also recapitulated a neuron-specific Ab response in 3/4 patients with established intra- cellular Ag-specific Abs (2 Hu+ AIE/PNS, 1 Ri+ AIE/PNS). In the patients who tested negative in the serum (no available CSF) with our CBA (AK5+ AIE), the antigenic target was detected with at a very low titer in validated diagnostic laboratories in both the serum and the CSF, a titer below the one we have set in our study. Indeed, when determining the level of dilution of IgG in serum and CSF, we deliberately selected a high dilution level to minimize false positive results and thus maximize the likelihood of detecting true positive ones. There were two additional patients with established AIE/PNS: 1 NMDAR and one PCA-Tr, whose antigens are expressed on the surface, respectively on the synapse, for whom we detected no reactive Abs either in the serum or the CSF. Indeed, our CBA entails permeabilization of the cells, which could potentially interfere with surface/synaptic Ag detection. These data suggest that our CBA assay is more able to detect intra-cellular than surface Ags and would thus make it particularly suitable for the search of CNS-specific intracellular Ag, such as in paraneoplastic conditions. Additionally, our protocol to obtain hiPSC-derived neurons does not generate Purkinje cells, possibly preventing from the detection of PCA-Tr+ auto-Abs. Detection of surface or synaptic antigens may rely on live cell-based assay, requiring further refinements for future investigations.

Most interestingly, beyond seropositive AQP4+ NMO and definite AIE/PNS, we identified astrocyte- and neuron-reactive Abs in 14/89 patients. For these patients, CNS-specific Abs were not reported at the time of blood sampling yet, including nine IND and four NIND patients. Remarkably, one patient (NIND #3), who suffered from amyotrophic lateral sclerosis showed a distinct astrocyte-binding reactivity as compared to the other astrocyte-reactive NIND patient. This observation is supported by recent observations of neural auto-Abs in this pathology.^36,37^ The two other NIND patients exhibited a reactivity against neurons that was restricted to the serum, contrasting to what was observed in most of IND patients for whom both serum and CSF were detected positive when paired samples were tested. In line with the latter observations, using the CBA, the intensity of the CNS-reactive IgG response was more intense in IND than in NIND patients, either when it was directed against astrocytes (in serum the astrocyte-reactivity was present only in IND; CSF IND 2.47-fold higher than NIND) or against neurons (serum IND 1.25-fold higher than NIND; CSF IND 4.69-fold higher than NIND). Additionally, we found that the CSF is a better compartment than the serum to screen for CNS-reactive Ab. These findings support recent reports emphasizing the importance of CSF testing for accurate diagnostic interpretation, particularly neuronal Ab reactivity.^38–40^ Noteworthy, the patients for whom we newly identified astrocyte- and/or neuron-reactive patients with our 96-well hiPSC-derived CNS CBA, all had a negative result using mouse tissue as an antibody detection method in a reference lab. These findings thus support our human based model as a complementary tool for the identification of not yet reported CNS-reactive.

One could argue that this CBA is currently restricted to astrocytes and neurons. Including other CNS cell types such as oligodendrocytes, microglia, endothelial or even ependymal cells to broaden antigenic diversity would certainly be an asset. However, astrocytes and neurons currently cover the vast majority of the CNS Ags reported to date to be associated with neurological disorders.^41^ MOG is the only antigen not expressed by astrocytes and/or neurons, but by oligodendrocytes.^11^ Recently, authors have relied on new elegant peptide-based techniques such as phage-based techniques to detect CNS-reactive auto-Abs and their cognate Ags.^42–46^ Nevertheless, even though these proteome-wide auto-Ab discovery platforms are unbiased, these techniques are not ideal for detecting proteins in their 3D conformation as they do not consider post-translational protein modifications. Yet these parameters are crucial for Ab-antigen recognition.^47–49^ We think that our unique human hiPSC-based platform may serve as a rapid screening tool in patients with a clinical presentation suggestive for CNS-reactive Ab-mediated disease but in whom no anti-CNS cell Abs were detected in validated routine laboratory testing.

Beyond this screening, our platform could offer the possibility to identify previously unreported CNS Ags targeted by CNS-reactive Abs especially when considering intracellular Ag(s). Ultimately, our rapid 96-well hiPSC-derived CNS platform offers a promising avenue for further investigations ultimately translating into a better diagnosis and therapeutical management of auto-Ab-related neurological conditions.

## Acknowledgments

We thank G. Le Goff from the Neuroimmunology unit (CHUV, Lausanne, CH) for help in enrolling patients and obtaining blood samples; Lionel Arlettaz from the Wallis hospital (Sion, CH) for additional sample testing, Binxia Yang and Lilian Mills from the Mayo Clinic Neuroimmunology Laboratory (Rochester, US) for antibody testing.

## Funding

This work was supported by the Swiss National Science Foundation (320030_179531, CRSK- 3_190190), a generous donator advised by Carigest SA and the Fondation pour la médecine de Laboratoire F4LABMED.

## Disclosures

A. Mathias, S. Perriot, S. Jones, M. Canales, M. Gimenez, N. Torcida, L. Oberholster report no disclosures relevant to the manuscript.
R. Bernard-Valnet received travel grants from Roche and received speaker honoraria from Novartis. None were related to this work.
A. Hottinger has served as an expert in advisory boards from Novocure and Bayer, received speaker honoraria from Novocure, all paid to the institution and not related to this work.
A. Zekeridou has patents submitted for Tensacin-R IgG, PDE10A-IgG and DACH1-IgG as biomarkers of neurological autoimmunity. Receives research funding from Roche/Genetech non relevant to this work. Has consulted for Alexion Pharmaceutical without personal compensation.
M. Theaudin has served as an expert in advisory boards for Biogen, Genzyme-Sanofi, Merck, Novartis and Roche, received travel grants from Biogen, Genzyme-Sanofi, Merck, Novartis and Roche, received speaker honoraria from Biogen, Novartis and Merck. None were related to this work.
C. Pot has served as an expert in advisory boards and received travel grants from Biogen, Genzyme-Sanofi, Merck, Novartis and Roche none related to this work
R. Du Pasquier has served on scientific advisory boards for Biogen, BMS, Merck, Novartis, Roche, and Sanofi-Genzyme; has received funding for travel or speaker honoraria from Biogen, Merck, Roche, and Sanofi-Genzyme.

## Supplementary material

Supplementary material is available at BioRxiv online.

## Take-home points

- Novel cell-based assay using hiPSC-derived neurons and astrocytes.
- Detection of astrocyte- and neuronal-antibodies in 19 out of 99 subjects.
- Higher neural-reactive antibody binding observed in inflammatory neurological diseases.
- Promising tool for early diagnosis of immune-mediated neurological conditions.

## Notes

### Summary of Updates

updated table 1 and supplementary tables 3-7

