## Supplementary material for "Human stem cell derived neurons and astrocytes to detect novel auto-reactive IgG signature in immune-mediated neurological diseases"

#### Supplementary Figure Legends

**Supplementary Figure 1. Characterization of hiPSC-derived astrocytes and neurons used for the detection of CNS-reactive antibodies in the serum and CSF of selected patients.**

**(A)** The expression profiles of seven astrocytic, neuronal and hiPSC markers are regularly evaluated in hiPSC-derived astrocytes, neurons and NPCs from healthy donors used in this study (HD#002 and HD#003). One representative experiment is reported (LNISi002-B). The seven markers are listed on the y axis and the cell type and donor are listed on the x-axis. Results are expressed as the *Z-score* of the $-\Delta C_T$ ($C_T$ of gene of interest – $C_T$ of GAPDH).

**(B)** Representative immunofluorescence images of hiPSC-derived astrocytes expressing AQP4, CD44, GLAST, GFAP, S100β (red) obtained from healthy control HD#002.

**(C)** Representative immunofluorescence images of hiPSC-derived neurons expressing MAP2, β-Tubulin-III, HuC/D, NMDAR1 and NMDAR2B (red) obtained from HD#002.

**(B, C).** Nuclei stained in DAPI appear in blue. Control panel correspond to cells stained with secondary antibodies only. Scale bar, 50 µm.

**Supplementary Figure 2. Cross-validation of the automated plate reader rapid IgG measurement with single cell microscopy analyses.**

Human iPSC-derived astrocytes (A) and neurons (B) were exposed to serum or CSF. FI were assessed using a Synergy® microplate reader. Subsequent IgG indexes (*z-score*) were calculated (see Methods for details). The exact same wells of the CBA were then observed by fluorescence microscopy using a EVOS M700 automated microscope plate reader and at least four images were acquired per well. Single-cell associated IgG intensity were then measured in all images. Correlations between IgG indexes at the well level (*z-score*) [x-axis] and the median IgG intensity at the single cell level by microscopy (*z-score*) [y-axis] were represented using a generalized linear model (reference line in plain black, confidence interval set at 95% in shadowed area) and tested using a Spearman's rank correlation test (r and p values on the graphs).

**Supplementary Figure 3. Gating strategy of hiPSC-derived astrocytes and neurons vs PBMC and subsequent analyses of IgG binding to CNS cells vs CD3+ T cells.**

Cells (A. astrocytes, B. neurons, C. PBMC) were exposed to either IgG-detection antibodies only (Negative Control in grey), or selected patient CSF followed by IgG-detection Abs (A, C. GFAP+ astrocytopathy #1 in black, B-C. Ri+ PNS #1 in dark red). Total PBMCs were also stained using anti-CD3 antibodies. Dead cells were excluded using Live/dead marker. Representative gating strategies are represented. Data were acquired on LSRII flow and analyzed using Flow Jo Software.

Supplementary Figure 1.

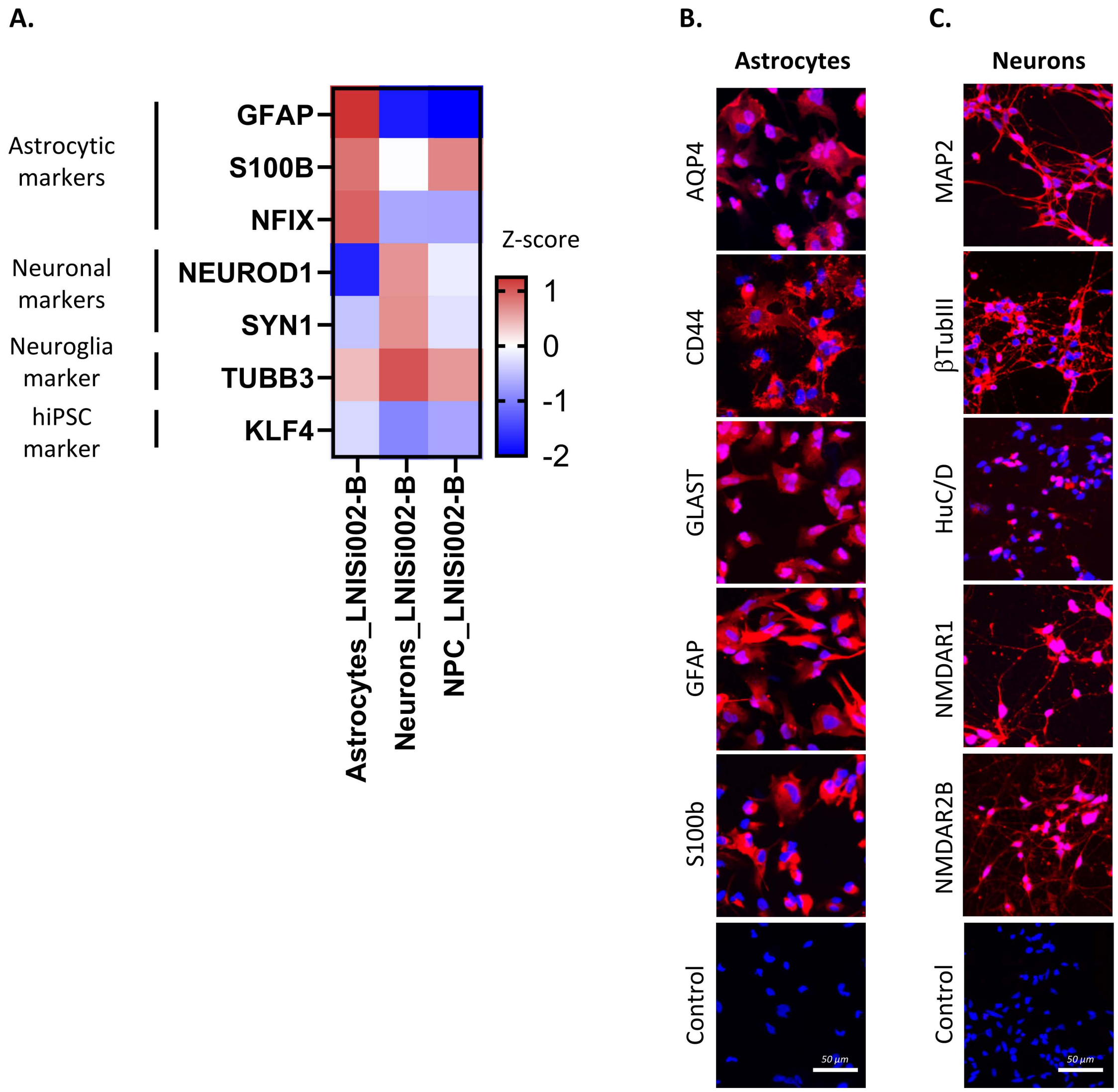

### Supplementary Figure 2.

**A.** Astrocytes – correlation plate reader : microscopy

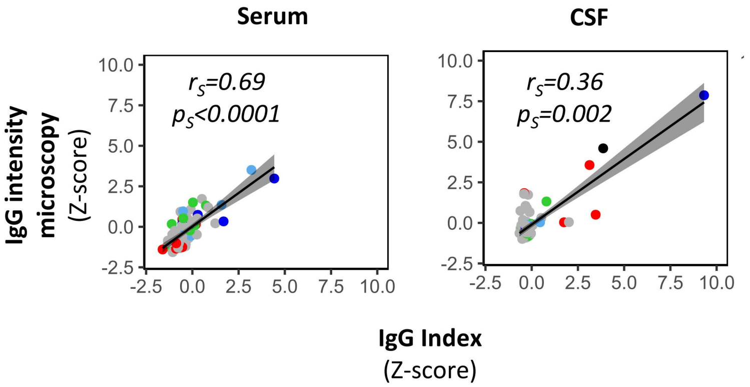


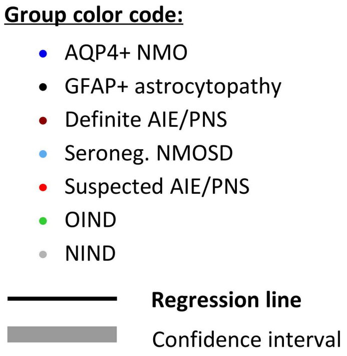


**B.** Neurons – correlation plate reader : microscopy

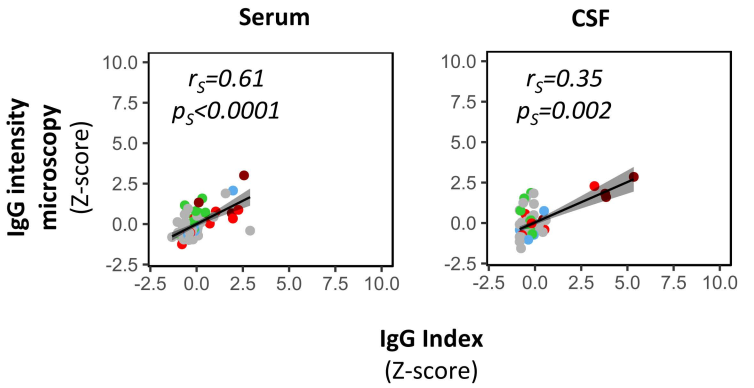

Supplementary Figure 3.

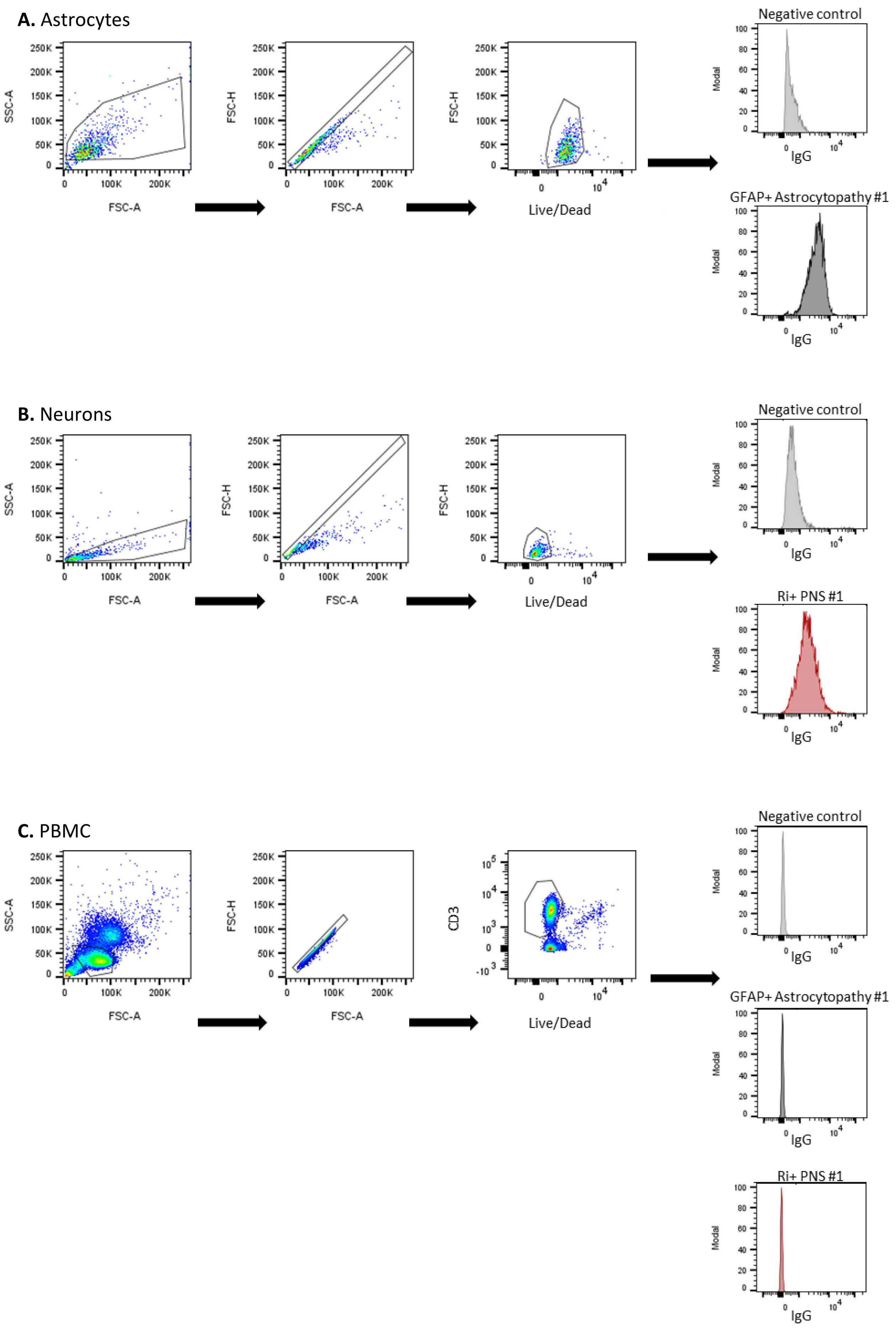

**Supplementary Table 1. List of primers used for reverse transcription quantitative PCR (RT-qPCR).**

| GENE NAME | FORWARD | REVERSE |
|---|---|---|
| **GFAP** | GCCAGTTGCAGTCCTTGAC | GCGCATCTGCCTCTCCA |
| **NFIX** | CAAGGAGATGCGGACATCAAAC | ACCCCGGAAGTCACAAAACA |
| **S100B** | GCAGCAAGGAGACCAGGAA | CCACCATGGCCTTCTCCA |
| **KLF4** | CTGCGGCAAAACCTACACAA | CGTCCCAGTCACAGTGGTAA |
| **NEUROD1** | GCCCCAGGGTTATGAGACTA | TCTGTCCAGCTTGGAGGAC |
| **SYN1** | GCAAGGACGGAAGGGATCA | TGTCTTCATCCTGGTGGTCAC |
| **TUBB3** | GAGCGGATCAGCGTCTACTA | GGTTCCAGGTCCACCAGAA |

**Supplementary Table 2. List of antibodies and detection reagents used for immunofluorescence assays.**

| Target | Clone | Manufacturer | Reference | Host | Dilution |
|---|---|---|---|---|---|
| ***Primary antibodies*** | | | | | |
| AQP4 | polyclonal | Life Technologies | PA5-53234 | rabbit | 1/50 |
| S100b | EP1576Y | Abcam | ab52642 | rabbit | 1/200 |
| GLAST | polyclonal | Abcam | ab416 | rabbit | 1/200 |
| GFAP | polyclonal | Millipore | Ab5804 | rabbit | 1/200 |
| CD44 | DB105 | Miltenyi | 130-113-333 | biot | 1/50 |
| NMDAR1 | polyclonal | Life Technologies | PA3-102 | rabbit | 1/50 |
| NMDAR2B | polyclonal | Life Technologies | PA3-105 | rabbit | 1/50 |
| btubIII | TuJ-1 | biotechne | BAM1195 | biot | 1/100 |
| HuC/HuD | 16A11 | Life Technologies | A-21271 | mouse | 1/50 |
| MAP2 | polyclonal | Abcam | AB5392 | chicken | 1/200 |
| ***Detection antibodies/reagents*** | | | | | |
| anti-rabbit AF546 | polyclonal | LIFE TECHNOLOGIES EUROPE BV | A10040 | donkey | 1/200 |
| anti-mouse AF546 | polyclonal | LIFE TECHNOLOGIES EUROPE BV | A10036 | donkey | 1/200 |
| anti-chicken AF546 | polyclonal | LIFE TECHNOLOGIES EUROPE BV | A11040 | goat | 1/200 |
| anti-human IgG-biotin | poylclonal | Invitrogen | 31774 | goat | 1/100 |
| strepatvidin-PE | -- | BD | 554061 | -- | 1/1000 |

**Supplementary Table 3. Individual clinical information of the AQP4+ NMO patients enrolled in this study.**

| Subject | Immune modifying treatment[a] | Treatment duration[a] | Tested sample(s) | Outlier 96-well CBA neuron[b] | Outlier 96-well CBA astrocyte[b] |
|---|---|---|---|---|---|
| AQP4+ NMO #1 | N/A | -- | SERUM/CSF | NO | **YES** (Serum/CSF) |
| AQP4+ NMO #2 | Rituximab | 39 days | SERUM | NO | **YES** (serum) |
| AQP4+ NMO #3 | Mycophenolate Mofetil | 13.7 years | SERUM/CSF | NO | NO |
| AQP4+ NMO #4 | Mycophenolate Mofetil | 7.1 years | SERUM | NO | NO |

[a] Times expressed at sampling date
[b] Results from the 96-well hiPSC-derived CNS CBA are expressed as outlier YES or NO. If YES, in brackets the sample(s) to be tested as an outlier in the corresponding 96-well hiPSC-derived CNS CBA either using neurons or astrocytes.
Abbreviations: AQP4+ NMO, AQP4+ neuromyelitis optica; CSF, cerebrospinal fluid; N/A, not applicable.

**Supplementary Table 4. Individual clinical information of the Seronegative NMOSD patients enrolled in this study.**

| Subject | Clinical manifestations | Immune modifying treatment[a] | Treatment duration[a] | Tested sample(s) | Outlier 96-well CBA neuron[b] | Outlier 96-well CBA astrocyte[b] | Mouse TIF evaluation[c] |
|---|---|---|---|---|---|---|---|
| Seronegative NMOSD #1 | Severe bilateral ON and area postrema syndrome (hiccups) | N/A | -- | SERUM/CSF | **YES** (Serum/CSF) | **YES** (Serum/CSF) | NO |
| Seronegative NMOSD #2 | myelitis and ON | N/A | -- | SERUM | NO | **YES** (Serum) | NO |
| Seronegative NMOSD #3 | myelitis and optic neuritis | N/A | -- | SERUM/CSF | NO | NO | NO |
| Seronegative NMOSD #4 | bilateral ON and myelitis | Azathioprin | 53 days | SERUM/CSF | NO | NO | NO |
| Seronegative NMOSD #5 | severe ON and myelitis | Rituximab | 0 days | SERUM | NO | NO | NO |
| Seronegative NMOSD #6 | severe bilateral ON and myelitis | Azathioprin | 2.3 years | SERUM/CSF | NO | NO | NO |

[a] Times expressed at sampling date

[b] Results from the 96-well hiPSC-derived CNS CBA are expressed as outlier YES or NO. If YES, in brackets the sample(s) to be tested as an outlier in the corresponding 96-well hiPSC-derived CNS CBA either using neurons or astrocytes.

[c] Samples from patients found to be positive in our novel 96-well hiPSC-derived CNS CBA were evaluated by indirect immunofluorescence on a mouse tissue composite (TIF) (YES or NO). If YES, in brackets the result of the mouse TIF.

Abbreviations: NMOSD : neuromyelitis optica spectrum disorder; ON, optic nerve; CSF, cerebrospinal fluid; N/A, not applicable.

**Supplementary Table 5. Individual clinical information of the definite AIE/PNS patients enrolled in this study.**

| Subject | Immune modifying treatment[a] | Treatment duration[a] | Tested sample(s) | Outlier 96-well CBA neuron[b] | Outlier 96-well CBA astrocyte[b] |
|---|---|---|---|---|---|
| anti-AK5+ AIE/PNS #1 | N/A | -- | SERUM | NO | NO |
| anti-Hu+ AIE/PNS #1 | Methylprednisolone | 12 days | SERUM/CSF | **YES** (Serum/CSF) | NO |
| anti-Hu+ AIE/PNS #2 | N/A | -- | CSF | **YES** (CSF) | NO |
| anti-NMDA-R+ AIE/PNS #1 | N/A | -- | SERUM/CSF | NO | NO |
| anti-PCA-Tr+ AIE/PNS #1 | N/A | -- | SERUM | NO | NO |
| anti-Ri+ AIE/PNS #1 | N/A | -- | SERUM/CSF | **YES** (Serum/CSF) | NO |
| anti-GFAP+ astrocytopathy #1 | N/A | -- | CSF | NO | **YES** (CSF) |

[a] Times expressed at sampling date
[b] Results from the 96-well hiPSC-derived CNS CBA are expressed as outlier YES or NO. If YES, in brackets the sample(s) to be tested as an outlier in the corresponding 96-well hiPSC-derived CNS CBA either using neurons or astrocytes.
Abbreviations: AIE : autoimmune encephalitis; PNS : paraneoplastic syndrome; CSF, cerebrospinal fluid; N/A, not applicable.

**Supplementary Table 6. Individual clinical information of the Suspected AIE/PNS patients enrolled in this study.**

| Subject | Clinical manifestations | Tumor/oncologic condition | Immune modifying treatment[a] | Treatment duration[a] | Tested sample(s) | Outlier 96-well CBA neuron[b] | Outlier 96-well CBA astrocyte[b] | Mouse TIF evaluation[c] |
|---|---|---|---|---|---|---|---|---|
| Suspected AIE/PNS #1 | opsoclonus myoclonus | N/A | N/A | -- | CSF | NO | **YES** (CSF) | **YES** (neg) |
| Suspected AIE/PNS #2 | limbic encephalitis | N/A | N/A | -- | SERUM/CSF | **YES** (Serum) | NO | NO |
| Suspected AIE/PNS #3 | limbic encephalitis | kidney tumor | N/A | -- | SERUM/CSF | NO | NO | NO |
| Suspected AIE/PNS #4 | encephalomyelitis | myeloid leukemia | N/A | -- | CSF | **YES** (CSF) | NO | NO |
| Suspected AIE/PNS #5 | encephalomyelitis | B cell lymphoma | IFN-b1a | 14 days | CSF | NO | **YES** (CSF) | **YES** (neg) |
| Suspected AIE/PNS #6 | Myasthenia gravis | thymoma | N/A | -- | SERUM | NO | NO | NO |
| Suspected AIE/PNS #7 | post-ICI oculomotor nerves myositis | melanoma with lung metastasis | pembrolizumab --> ipilimumab --> nivolumumab | 3.3 months | SERUM | NO | NO | NO |
| Suspected AIE/PNS #8 | opsoclonus myoclonus; cerebellar ataxia; dementia | N/A | N/A | -- | SERUM | NO | NO | NO |
| Suspected AIE/PNS #9 | limbic encephalitis, epilepsy, catatonia | neuroendocrine tumor (para-trackea) | N/A | -- | CSF | **YES** (CSF) | **YES** (CSF) | **YES** (neg) |
| Suspected AIE/PNS #10 | Cerebellar degeneration | ovarian adenocarcinoma | Prednisone | 2 months | SERUM | NO | NO | NO |
| Suspected AIE/PNS #11 | Stiff person syndrome | neuroendocrine tumor (ileum) | N/A | -- | SERUM | **YES** (Serum) | NO | YES |
| Suspected AIE/PNS #12 | opsoclonus myoclonus | ovarian teratoma | N/A | -- | SERUM/CSF | NO | NO | NO |

[a] Times expressed at sampling date
[b] Results from the 96-well hiPSC-derived CNS CBA are expressed as outlier YES or NO. If YES, in brackets the sample(s) to be tested as an outlier in the corresponding 96-well hiPSC-derived CNS CBA either using neurons or astrocytes.

[c] Samples from patients found to be positive in our novel 96-well hiPSC-derived CNS CBA were evaluated by indirect immunofluorescence on a mouse tissue composite (TIF) (YES or NO). If YES, in brackets the result of the mouse TIF.
Abbreviations: AIE : autoimmune encephalitis; PNS : paraneoplastic syndrome; N/A, not applicable ; CSF, cerebrospinal fluid; neg, negative.

**Supplementary Table 7. Individual clinical information of the OIND and NIND patients enrolled in this study.**

| Subject | Diagnostic | Immune modifying treatment[a] | Treatment duration[a] | Tested sample(s) | Outlier 96-well CBA neuron[b] | Outlier 96-well CBA astrocyte[b] | Mouse TIF evaluation[c] |
|---|---|---|---|---|---|---|---|
| OIND #1 | Inflammatory temporal lobe mass | N/A | -- | SERUM/CSF | NO | **YES** (Serum/CSF) | **YES** (Neg) |
| OIND #2 | Behcet Disease | Prednisone | 53 days | SERUM/CSF | NO | NO | NO |
| OIND #3 | Chronic meningitis (unknow origin) | N/A | -- | SERUM | NO | NO | NO |
| OIND #4 | Inflammatory cranial neuritis | N/A | -- | SERUM | NO | NO | NO |
| OIND #5 | Myelitis of unknown origin | N/A | -- | CSF | NO | NO | NO |
| OIND #6 | Susac syndrome | Rituximab | 9.3 months | SERUM | NO | NO | NO |
| OIND #7 | Susac syndrome | N/A | -- | SERUM/CSF | NO | NO | NO |
| OIND #8 | Unknown demyelinating disease | N/A | -- | SERUM/CSF | NO | NO | NO |
| OIND #9 | Vasculitis | N/A | -- | CSF | NO | NO | NO |
| OIND #10 | Vasculitis | N/A | -- | SERUM | NO | NO | NO |
| OIND #11 | Vasculitis | N/A | -- | SERUM/CSF | NO | NO | NO |
| OIND #12 | Vogt-Koyanagi-Harada syndrome | N/A | -- | SERUM/CSF | NO | NO | NO |
| OIND #13 | Mutlifocal mononeuropathy | N/A | -- | SERUM/CSF | NO | NO | NO |
| NIND #1 | Acrocyanosis | N/A | -- | SERUM/CSF | NO | NO | NO |
| NIND #2 | Acroparesthesia | N/A | -- | SERUM/CSF | NO | NO | NO |
| NIND #3 | Amyotrophic lateral sclerosis | N/A | -- | CSF | NO | **YES** (CSF) | **YES** (Neg) |
| NIND #4 | Amyotrophic lateral sclerosis | N/A | -- | SERUM/CSF | NO | NO | NO |
| NIND #5 | Amyotrophic lateral sclerosis | N/A | -- | CSF | NO | NO | NO |
| NIND #6 | Amyotrophic lateral sclerosis | N/A | -- | CSF | NO | NO | NO |
| NIND #7 | Amyotrophic lateral sclerosis | N/A | -- | CSF | NO | NO | NO |
| NIND #8 | Amyotrophic lateral sclerosis | N/A | -- | CSF | NO | NO | NO |
| NIND #9 | Catatonia with unknown origin | N/A | -- | SERUM/CSF | NO | NO | NO |
| NIND #10 | Cognitive disorders (memory and attention deficits | N/A | -- | SERUM/CSF | NO | NO | NO |
| NIND #11 | Depression | N/A | -- | SERUM/CSF | NO | NO | NO |
| NIND #12 | Depression | N/A | -- | SERUM/CSF | NO | NO | NO |
| NIND #13 | Depression | N/A | -- | SERUM | NO | NO | NO |
| NIND #14 | Dysesthesias | N/A | -- | SERUM/CSF | NO | NO | NO |

| | | | | | | | |
|---|---|---|---|---|---|---|---|
| NIND #15 | Dysesthesias | N/A | -- | SERUM/CSF | NO | NO | NO |
| NIND #16 | Epilepsy | N/A | -- | SERUM/CSF | NO | NO | NO |
| NIND #17 | Epilepsy | N/A | -- | CSF | NO | NO | NO |
| NIND #18 | Fibromyalgia | N/A | -- | CSF | NO | NO | NO |
| NIND #19 | Functional syndrome | N/A | -- | SERUM/CSF | NO | NO | NO |
| NIND #20 | Genetic myelopathy | N/A | -- | SERUM/CSF | NO | NO | NO |
| NIND #21 | Headache | N/A | -- | CSF | NO | NO | NO |
| NIND #22 | Headache | N/A | -- | CSF | NO | NO | NO |
| NIND #23 | Inclusion-body myositis | N/A | -- | CSF | NO | **YES** (CSF) | **YES** (Neg) |
| NIND #24 | idiopathic facial palsy | N/A | -- | SERUM/CSF | NO | NO | NO |
| NIND #25 | Intracranial hypertension | N/A | -- | SERUM/CSF | NO | NO | NO |
| NIND #26 | Intracranial hypertension | N/A | -- | SERUM/CSF | NO | NO | NO |
| NIND #27 | Intracranial hypertension | N/A | -- | CSF | NO | NO | NO |
| NIND #28 | Ischaemic myelopathy | N/A | -- | SERUM/CSF | NO | NO | NO |
| NIND #29 | Kleine Levin Syndrome | N/A | -- | SERUM | NO | NO | NO |
| NIND #30 | Migraine | N/A | -- | SERUM | NO | NO | NO |
| NIND #31 | Migraine | N/A | -- | SERUM/CSF | NO | NO | NO |
| NIND #32 | Migraine | N/A | -- | SERUM/CSF | NO | NO | NO |
| NIND #33 | Migraine | N/A | -- | SERUM/CSF | NO | NO | NO |
| NIND #34 | Migraine | N/A | -- | SERUM/CSF | NO | NO | NO |
| NIND #35 | Migraine | N/A | -- | SERUM | NO | NO | NO |
| NIND #36 | Mononeuropathy | N/A | -- | SERUM/CSF | NO | NO | NO |
| NIND #37 | Non ischemic cerebral enhancing | N/A | -- | SERUM/CSF | **YES** (Serum) | NO | **YES** (Neg) |
| NIND #38 | non inflammatory CNS white matter lesions | N/A | -- | SERUM/CSF | NO | NO | NO |
| NIND #39 | optic chiasma syndrome | N/A | -- | SERUM/CSF | NO | NO | NO |
| NIND #40 | Parkinsonism syndrome | N/A | -- | CSF | NO | NO | NO |
| NIND #41 | Parkinsonism syndrome | N/A | -- | SERUM/CSF | NO | NO | NO |
| NIND #42 | Parkinsonism syndrome | N/A | -- | SERUM/CSF | NO | NO | NO |
| NIND #43 | Parkinsonism syndrome | N/A | -- | SERUM/CSF | NO | NO | NO |
| NIND #44 | Parkinsonism syndrome | N/A | -- | CSF | NO | NO | NO |
| NIND #45 | Parkinsonism syndrome | N/A | -- | CSF | NO | NO | NO |

| | | | | | | | |
|---|---|---|---|---|---|---|---|
| NIND #46 | Perinatal periventricular lesion | N/A | -- | SERUM/CSF | NO | NO | NO |
| NIND #47 | Radiculopathy | N/A | -- | SERUM/CSF | NO | NO | NO |
| NIND #48 | Radiculopathy | N/A | -- | SERUM/CSF | NO | NO | NO |
| NIND #49 | Radiculopathy | N/A | -- | SERUM/CSF | NO | NO | NO |
| NIND #50 | Reversible cerebral vasoconstriction syndrome | N/A | -- | SERUM/CSF | NO | NO | NO |
| NIND #51 | Spastic paraparesis | N/A | -- | SERUM/CSF | NO | NO | NO |
| NIND #52 | Spinal stenosis | N/A | -- | SERUM/CSF | NO | NO | NO |
| NIND #53 | Spinal stenosis | N/A | -- | CSF | NO | NO | NO |
| NIND #54 | Spinal stenosis | N/A | -- | CSF | NO | NO | NO |
| NIND #55 | Stroke | N/A | -- | SERUM/CSF | **YES** (Serum) | NO | NO |
| NIND #56 | Stroke | N/A | -- | SERUM/CSF | NO | NO | NO |
| NIND #57 | Stroke | N/A | -- | SERUM/CSF | NO | NO | NO |

[a] Times expressed at sampling date
[b] Results from the 96-well hiPSC-derived CNS CBA are expressed as outlier YES or NO. If YES, in brackets the sample(s) to be tested as an outlier in the corresponding 96-well hiPSC-derived CNS CBA either using neurons or astrocytes.
[c] Samples from patients found to be positive in our novel 96-well hiPSC-derived CNS CBA were evaluated by indirect immunofluorescence on a mouse tissue composite (TIF) (YES or NO). If YES, in brackets the result of the mouse TIF.
Abbreviations: OIND; other inflammatory neurological disorders; NIND, non-inflammatory neurological disorders; CSF, cerebrospinal fluid; N/A, not applicable.
